# Atomic insights into the signaling landscape of *E. coli* PhoQ Histidine Kinase from Molecular Dynamics simulations

**DOI:** 10.1101/2024.04.19.590235

**Authors:** Symela Lazaridi, Jing Yuan, Thomas Lemmin

## Abstract

Bacteria rely on two-component systems to sense environmental cues and regulate gene expression for adaptation. The PhoQ/PhoP system exemplifies this crucial role, playing a key part in sensing magnesium (Mg^2+^) levels, antimicrobial peptides, mild acidic pH, osmotic upshift, and long-chain unsaturated fatty acids, promoting virulence in certain bacterial species. However, the precise details of PhoQ activation remain elusive. To elucidate PhoQ’s signaling mechanism at atomic resolution, we combined AlphaFold2 predictions with molecular modeling and carried out extensive Molecular Dynamics (MD) simulations. Our MD simulations revealed three distinct PhoQ conformations that were validated by experimental data. Notably, one conformation was characterized by Mg^2+^ bridging the acidic patch in the sensor domain to the membrane, potentially representing a repressed state. Furthermore, the high hydration observed in a putative intermediate state lends support to the hypothesis of water-mediated conformational changes during PhoQ signaling. Our findings not only revealed specific conformations within the PhoQ signaling pathway, but also hold significant promise for understanding the broader histidine kinase family due to their shared structural features. Our approach paves the way for a more comprehensive understanding of histidine kinase signaling mechanisms across various bacterial species and opens the door for developing novel therapeutics that target PhoQ modulation.

## Introduction

Many microorganisms, encompassing bacteria, archaea, and fungi exhibit the remarkable ability to thrive in diverse and often challenging environments [1] [2]. They have evolved intricate regulatory systems to sense and respond to changes in their surroundings [3]. Among these regulatory mechanisms, two-component systems (TCSs) stand as fundamental complexes that orchestrate cellular adaptation [4] [5]. TCSs are composed of two distinct units: the histidine kinase (HK) and the response regulator (RR) proteins [6] [7]. HK acts as a sensor detecting a large spectrum of environmental stimuli, ranging from shifts in temperature or pH, to availability of nutrients or detection of antibiotics [8] [9] [10]. Upon activation, HK undergoes autophosphorylation at a highly conserved histidine residue, marking the onset of a sophisticated phosphorylation relay. This initiates a subsequent interaction between a HK and its cognate RR protein, leading to the transfer of the phosphoryl group typically onto an aspartate residue in the receiver domain of the RR. Finally, the phosphorylated RR dimerizes and binds to specific DNA sequences, regulating gene expression in response to the environmental signal [11] [12] [13]. The absence of TCSs from mammals makes them enticing targets for drug development [14].

PhoQ functions as the histidine kinase within the PhoQ/PhoP two-component system, notably abundant among gram-negative bacteria [15] [16]. Activation of PhoQ is initiated by a variety of environmental changes [17] [18], including a low concentration of Mg^2+^, acidic pH [19] [20], the presence of cationic antimicrobial peptides [21] [22] [23], osmotic upshift, and long-chain unsaturated fatty acids in bile. PhoP directly controls the transcription of an expansive array of genes, coordinating responses to environmental stress, alterations in the bacterial envelope’s composition, and modulating virulence [24] [25]. Structurally, PhoQ assembles into a transmembrane dimer with a molecular weight of about 110 kDa. Its configurations closely resemble the architecture of prototypical histidine kinases, comprised of five distinct domains: (i) a periplasmic sensor domain, (ii) a transmembrane (TM) domain linking the sensor domain to the cytosolic domains, which include (iii) the histidine kinases, adenylate cyclases, methyl-accepting chemotaxis proteins, and phosphatases (HAMP) domain [26], (iv) the dimerisation and histidine phosphotransfer (DHp) domain, and (v) the catalytic domain, also known as the ATP binding domain (Figure 1A) [27].

**Figure 1:**
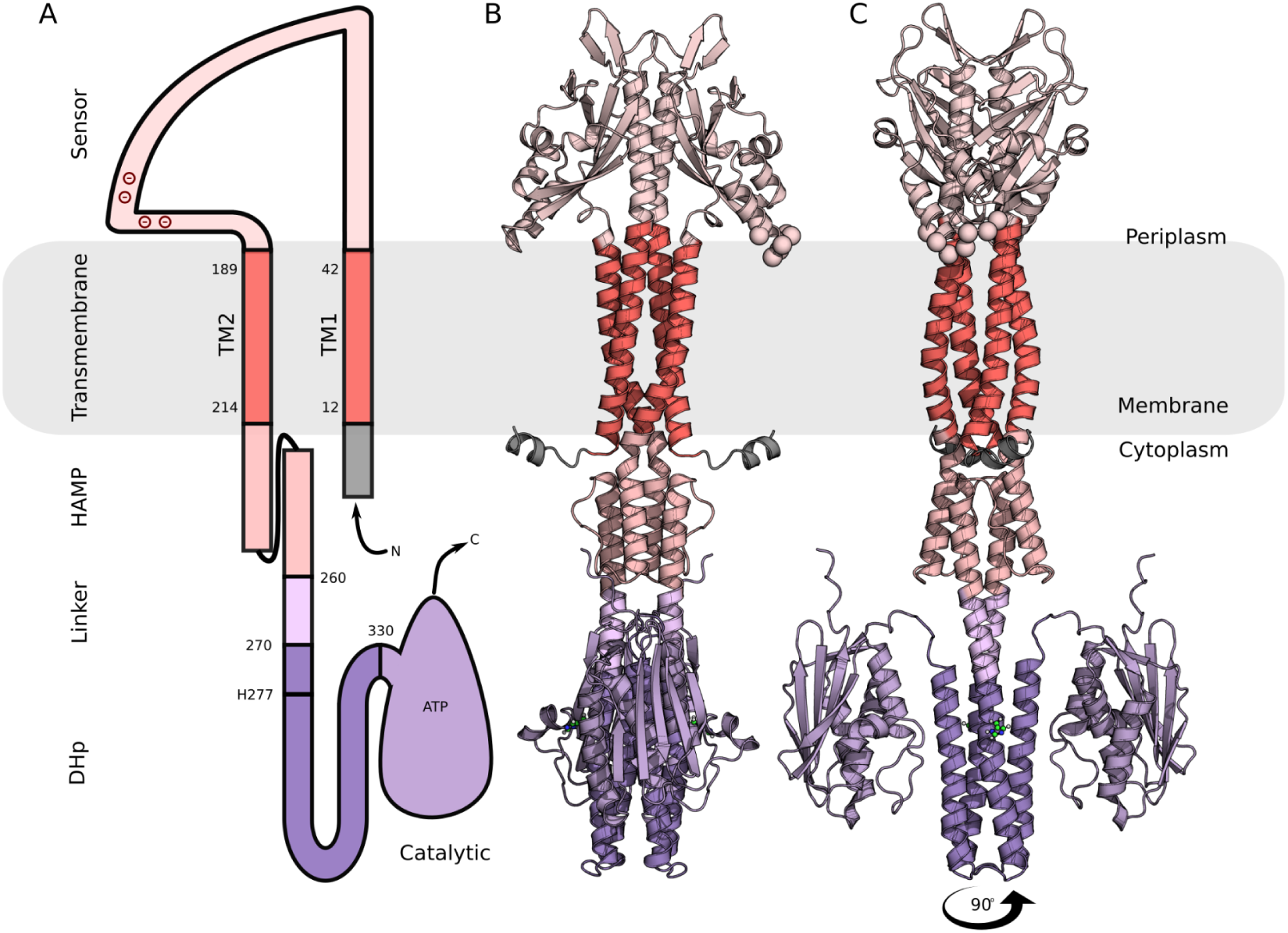
Predicted Structure of E. coli PhoQ. (A) Schematic representation of PhoQ topology. (B) Cartoon representation of the predicted PhoQ structure using AlphaFold2. Transmembrane domains (red), sensor and HAMP domains (shades of pink), and DhP and catalytic domains (shades of purple) are indicated. The membrane is depicted as a gray rectangle.

Several domains of PhoQ or homologous HKs have been experimentally characterized, providing valuable structural insights into components of the signaling mechanism. The sensor domain structure has been determined in both its monomeric and various oligomeric states [28], [29], [30], [31]. Through a combination of disulfide screen experiments and computational modeling, the dimeric form observed for *E. coli* (PDB id: 3bq8 [31]) has been suggested as the most likely representative of PhoQ’s physiologically relevant active conformation [32]. The asymmetry observed in this dimeric structure is believed to be essential for signaling, since autophosphorylation selectively takes place in one of the two subunits under physiological conditions [33], [34]. A proposed mechanism suggests that upon sensing stimuli, the sensor domain initiates structural rearrangements in the transmembrane domain, which are then propagated to the cytoplasmic domains [33].

In the absence of a high-resolution experimental structure for the TM domain, the phototaxis sensory rhodopsin II-transducer (HtrII, PDB id: 1h2s) complex [35] frequently serves as a model for HK transmembrane four-helical bundles [33], [36]. The PhoQ transmembrane domain harbors at its center a critical polar residue (N202) that likely forms a water cavity, similar to that observed in HtrII [37]. This cavity is believed to facilitate the arrangement of side chains around it, enabling the TM helix to bend and change conformation during signaling [35],[36]. This hypothesis is supported by ligand-dependent disulfide crosslinking experiments and Bayesian modeling, which identified two distinct structural states differing in the diagonal displacement of the TM helices [38], [39].

The HAMP [40] and DHp in the cytoplasm are well-conserved signal-transducing domains that have been extensively characterized experimentally in isolation or in combination with other cytoplasmic domains [41], [42], [43], [44]. Various signaling mechanisms have been proposed for the HAMP domain, including the gearbox [45] or the diagonal scissoring models [46]. The signaling initiated by the HAMP domain undergoes a transformation into an asymmetric conformation of the DHp bundle, mediated by a small connecting helix (S-helix). This distortion in the helical bundles of DHp facilitates the release of the catalytic domain, ultimately leading to the autophosphorylation of the conserved histidine [33], [41], [44].

Although previous studies have yielded crucial insights into the signal mechanism of PhoQ, a holistic understanding remains elusive due to the lack of comprehensive structural information for all the involved constituents. In this study, we therefore leveraged Alphafold2-multimer [47] to produce an atomic model of the complete PhoQ dimer. The predicted model exhibited strong congruence with existing structural data. To delve deeper into the intricate signaling mechanisms governing PhoQ, we combined classical molecular modeling and dynamics simulations and were able to capture two distinct conformations. These models were supported by available experimental data and could be potentially representative of an active and inactive conformation of PhoQ. Collectively, our findings provide new insights into signaling intricacies underpinning PhoQ, and our approach could be applicable to deciphering the signaling mechanisms of other histidine kinases.

## Results

### Structural characterization of AlphaFold prediction

AlphaFold2-multimer predicted with high confidence the full-length dimeric structure of *E. coli* PhoQ, achieving an average predicted local distance difference test (pLDDT) score of 0.8. [47] (Supplementary Figure S1 A). The generated model formed a symmetrical dimer, with a root mean square deviation (RMSD) of only 0.75 Å between the two monomers. The dimeric interface was characterized by a continuous helix (Figure 1B, C). Notably, there were marked drops in the pLDDT score in regions aligned well with the expected boundaries between the different domains of PhoQ (Figure S1 B). These discernible drops were leveraged to demarcate the start and end points of each domain for subsequent analyses.

The predicted PAS domain structure closely resembled the known crystallographic structure of *E. coli* (PDB ID: 3bq8) at the monomeric level, with a low RMSD of 0.6 Å [31]. However, significant deviations emerged in the predicted dimer (RMSD: 5.25 Å, Supplementary Figure S2). The interfacial helices in the model formed a shallower angle (10°) compared to the experimental structure (41°), impacting the orientation of the critical acidic patch (residues D136-D150) [48]. To quantify this difference, we embedded the Alphafold model into a lipid bilayer, using the orientations of proteins in membranes (OPM) server [49], and measured angles of 50.21° and 45.25° between the membrane surface and the acidic patch helices for chains A and B, respectively. To compare the orientations of the acidic patches in the experimental structure, we aligned the sensor interfaces (residues T48-K64) with the oriented Alphafold model. The angle for chain A remained comparable (47.02°), but was more parallel for chain B (25.85°), due to the inherent asymmetry observed in the experimental structure. Additionally, the crucial salt bridge between Asp179 and Arg50’ [31] present in the experimental structure was absent in the prediction, despite a comparable Cα-Cα distance (approximately 11.4 Å for both).

The transmembrane domain of PhoQ formed a four-helical bundle with antiparallel helices per monomer, defined as TM1 (S12 - V42) and TM2 (Y189 - A214). This structural arrangement is strongly corroborated by prior research [31], [32], [50]. Both TM1–TM1’ and TM2–TM2’ assembled in coiled coils with single-residue insertions disrupting the typical helical packing at positions V25 and S200, respectively (accommodation index of 1.0 [51]). To assess the novelty of this fold, we employed the FoldSeek search tool to query the Protein Data Bank (PDB) (Table S1) [52]. The four-helix bundle aligned with various structures; however few were transmembrane proteins (best PDB ID: 6sss_C, TM-score: 0.46, RMSD: 3.90 Å). This probably reflects the scarcity of such structures in the database. In contrast, comparison with the commonly used HTrII transmembrane bundle [36] using the TM-align mode [53] (RCSB.org [54]), yielded a significantly higher TM-score of 0.55 and a reduced RMSD of 2.56 Å. Additionally, we compared the Alphafold model to a previously published atomic model of the transmembrane domain [39] (Supplementary Figure S3A, B). Despite some variations (increased TM1 separation and closer TM2 helices in the published model, RMSD: 3.8 Å, TM-score: 0.56), the overall folds aligned well, enabling a seamless connection between the sensor interface helices and the HAMP domain (Supplementary Figure S3C).

The predicted HAMP domain adopted the expected two-antiparallel helix configuration, connected by a 10-residue loop. The FoldSeek search against the PDB database identified three HAMP domains from sensor proteins among the top hits but with low structural similarity (TM-Score: 0.6). The key difference resided in the interhelical angle. This variability was further confirmed by a statistical analysis (principal components analysis) of the identified HAMP domains, which revealed a wide distribution of angles, suggesting a lack of a single preferred conformation for the HAMP domain in these related sensor proteins.

The predicted DHp domain consisted of two alpha helices (a1 and a2) connected by a hairpin loop, forming a left-handed four-helix bundle in the dimer. This structure closely resembles other experimentally determined DHp domains (best TM-score: 0.84). Notably, the conserved histidine residue (H277), critical for the phosphorylation cascade, resides within the a1 helix with its side chain exposed to the cytoplasm. Moreover, the handedness of the bundle places the intra-protomeric catalytic domain in close proximity to this conserved histidine, strongly suggesting a cis-autophosphorylation mechanism [55].

The catalytic domain adopted the expected Bergerat fold and aligns, closely matching the experimentally solved structure of *E. coli* (PDB id: 1id0, RMSD=1.7Å) [56]. However, a key structural difference resided in the loop forming the ATP-lid, which connects helix α3 and beta-strand β3. In the 1id0 structure, the ATP-lid extended further, exposing the ligand to the solvent (Supplementary Figure S4). Although Alphafold was not trained with ligands, it has been shown that the configuration of binding pockets can often accommodate them [57]. We successfully positioned the ATP molecule and Mg^2+^ cation from the experimental structure (1id0) into the predicted binding pocket by aligning the main helices of the catalytic domains. In both structures, the Mg^2+^ cation was chelated by three oxygens of the triphosphate head and an additional interaction with N388 or N389, N385, and Q442 in the predicted and experimental structures, respectively. Conserved interactions were observed between K392 and N389 with the triphosphate head in both structures. However, the predicted structure lacks interactions with the ATP-lid residues R434 and R439. Furthermore, the adenosine moiety of the ATP engaged in pi-pi interactions with Y393 and formed a hydrogen bond with D415 in both structures

### Structural refinement with all-atom MD simulations

In order to further refine the predicted structure of the full-length PhoQ dimer, we carried out a set of all-atom molecular dynamics simulations. The AlphaFold model (*PhoQ_AF_*) was inserted into a bacterial mimic membrane bilayer and three replicas of all-atom MD simulations were conducted. These simulations were performed under near-physiological conditions (310K and 1 atm pressure), with each replica extending up to approximately 1.4 microseconds (Supplementary Table S2). The models remained stable throughout the simulations (average RMSD: 8.76 ± 2.57 Å, Supplementary Figure S5). The major difference was associated with the rearrangement of the catalytic domains, moving slightly away from the DHp. The protein’s core exhibited only marginal deviation from the predicted model, maintaining an average RMSD of 4.23 ± 1.28 Å (Figure 2A and Supplementary Figure S6A). The cross angle between the helices of the sensor interface remained consistently stable, averaging 7.38° ± 3.76° (Supplementary Figure S7). During the simulations, we also observed the dynamic formation of a salt bridge between D179 and R50’ (42% of simulation time with a cut-off distance of 3.6 Å) and R50 and D179’ (22% of simulation time).

**Figure 2:**
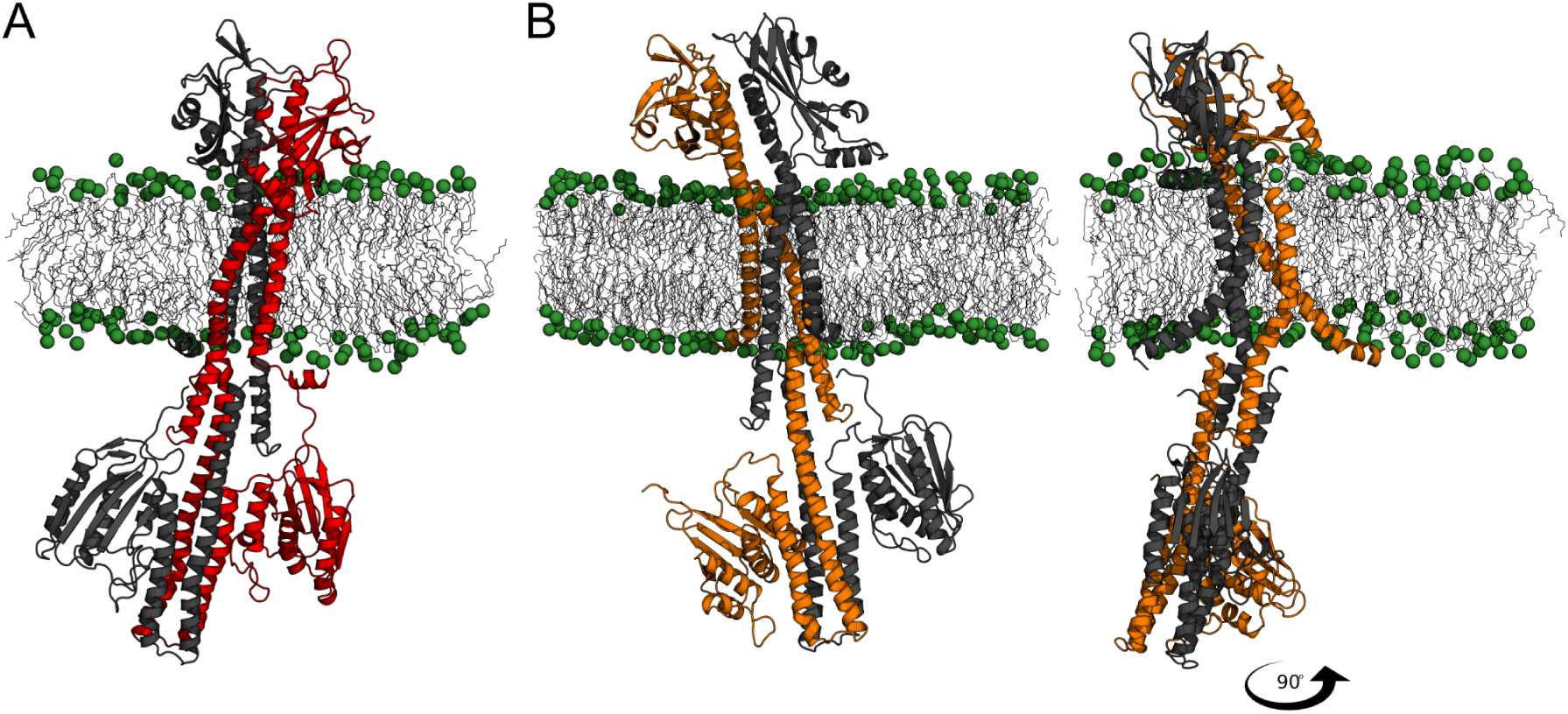
Representative structural models of PhoQ after MD simulation refinement. Cartoon representations of (A) *PhoQ_AF_* and (B) *PhoQ^TMD^_AF_* at the end of the MD simulation. The different chains are colored in gray and red or orange, respectively. The hydrophobic core of the membrane is depicted with gray sticks, and phosphate groups are shown as green spheres.

We attempted to refine the AlphaFold model’s sensor interface using targeted molecular dynamics (TMD) simulations (*PhoQ^TMD^_AF_*) to match the conformation observed in the experimental dimer structure (PDB ID: 3bq8). However, this introduced significant structural distortions (RMSD of full protein: 10.32 ± 3.76Å and core: 5.19 ± 2.03Å). In particular, the helix connecting the sensor to TM1 partially unfolded (residues D45 to T47), potentially leading to the decoupling of the sensor domain from the transmembrane region (Supplementary Figure S6B). We concluded that the TMD simulation employed was insufficient to induce the desired conformational alteration in the AlphaFold model, suggesting a more intricate structural coupling between the sensor and transmembrane domains that may hinder the desired conformational change in the model.

Given the good structural alignment between the previously modeled transmembrane [39] and *PhoQ^TMD^_AF_*, we constructed a hybrid model (*PhoQ^TMD^_AF_*) by combining the modeled transmembrane domain with the AlphaFold-predicted sensor and cytoplasmic domains (Supplementary Figure S6C). We then performed all-atom MD simulations in triplicate to refine the structure and assess its stability. The core of PhoQ remained stable throughout the simulations (4.16 ± 0.47Å, Supplementary Figure S5) and the sensor interfacial helices maintained a parallel alignment (cross angle: 13.10 ± 4.00°, Supplementary Figure S7). The simulations also captured the release of one of the catalytic domains.

Encouraged by the stability of the *PhoQ_H_*, we conducted a TMD simulation (*PhoQ^TMD^_H_*) to induce an interface angle resembling the experimental dimer (PDB ID: 3bq8). Even though the structure remained stable during the subsequent 1.5 µs unrestrained MD simulations, conducted in three replicas, (RMSD of 7.46 ± 3.24 Å, Supplementary Figure S5), the resulting conformation deviated significantly from *PhoQ_AF_* (Figure 2B and Supplementary Figure S6D). The equilibrated conformation of *PhoQ^TMD^_H_* was characterized by a highly asymmetric and curved structure, with several kinks along the core. Notably, the sensor domain tilted significantly (49.12° ± 5.11°) relative to the membrane compared to its perpendicular orientation in *PhoQ^TMD^_AF_*. Furthermore, a kink at position F44 in chain B allowed the sensor interface to retain a cross-angle more closely aligned with the experimental structure, approximately 27.22° ± 4.63°. This conformation was similar to the experimental structure (PDB ID: 3bq8, RMSD: 4.07 ± 0.44 Å). The interface helices pivoted around E55, while the E55 (chain A) and R53 (chain B) formed a distance of 3.7 Å for 33% of the simulation time, agreeing with the experimental structure. The increased cross-angle led to a near vertical alignment of the acidic patch helix on chain B (∼ 71.2°), bringing the acidic cluster close to the membrane surface. In contrast, the acidic patch helix of the other protomer aligned almost parallel to the membrane surface, forming an angle of approximately 7°. Although the conformations of the HAMP and DHP domains remained very similar to *PhoQ_AF_* (1.82 ± 0.30 Å RMSD), the catalytic domain of chain A was released and pivoted towards the DHp region.

### Conformational change of the transmembrane bundle

To understand how the altered transmembrane domain between *PhoQ_AF_* and *PhoQ^TMD^_H_* affected the protein structure, we investigated structural deviations propagated along this domain. We divided the transmembrane domain into three slices, each defined by two alpha carbons per protomer and oriented parallel to the membrane (Figure 3 and Supplementary Figure S8). The top slice (closest to the sensor domain) comprised residues A36 and W194, the middle slice (center of the membrane) residues L28 and N202, and the bottom slice (closest to the HAMP domain) residues T21 and L210. Compared to *PhoQ^TMD^_AF_*, *PhoQ^TMD^_H_* exhibited consistently larger TM1-TM1’ distances and smaller TM2-TM2’ distances. This resulted in TM1-TM1’ helices forming a cone shape, narrowing towards the sensor domain, whereas TM2-TM2’ helices adopted an inverted cone shape. Interestingly, despite the overall asymmetry of *PhoQ^TMD^_H_*, the arrangement of residues within each slice displayed a more symmetrical, diamond-like shape compared to the distorted appearance in *PhoQ_AF_* (Figures 3 and Supplementary Figure S8). The configuration of *PhoQ^TMD^_H_* was accompanied by a bent at I207 of approximately ∼3.5 degrees proximal to P208 in chain A (compared to ∼6.8 degrees in *PhoQ_AF_*), and a pistonning movement between transmembrane helices (Supplementary Figure S9). A partial unfolding on the connecting region between TM2 and the sensor domain (I181 – Y189) was also observed in *PhoQ^TMD^_H_*. In both models, the amphipathic N-terminal helices of TM1 (M1 to P8) were folded back and bound to the membrane surface.

**Figure 3:**
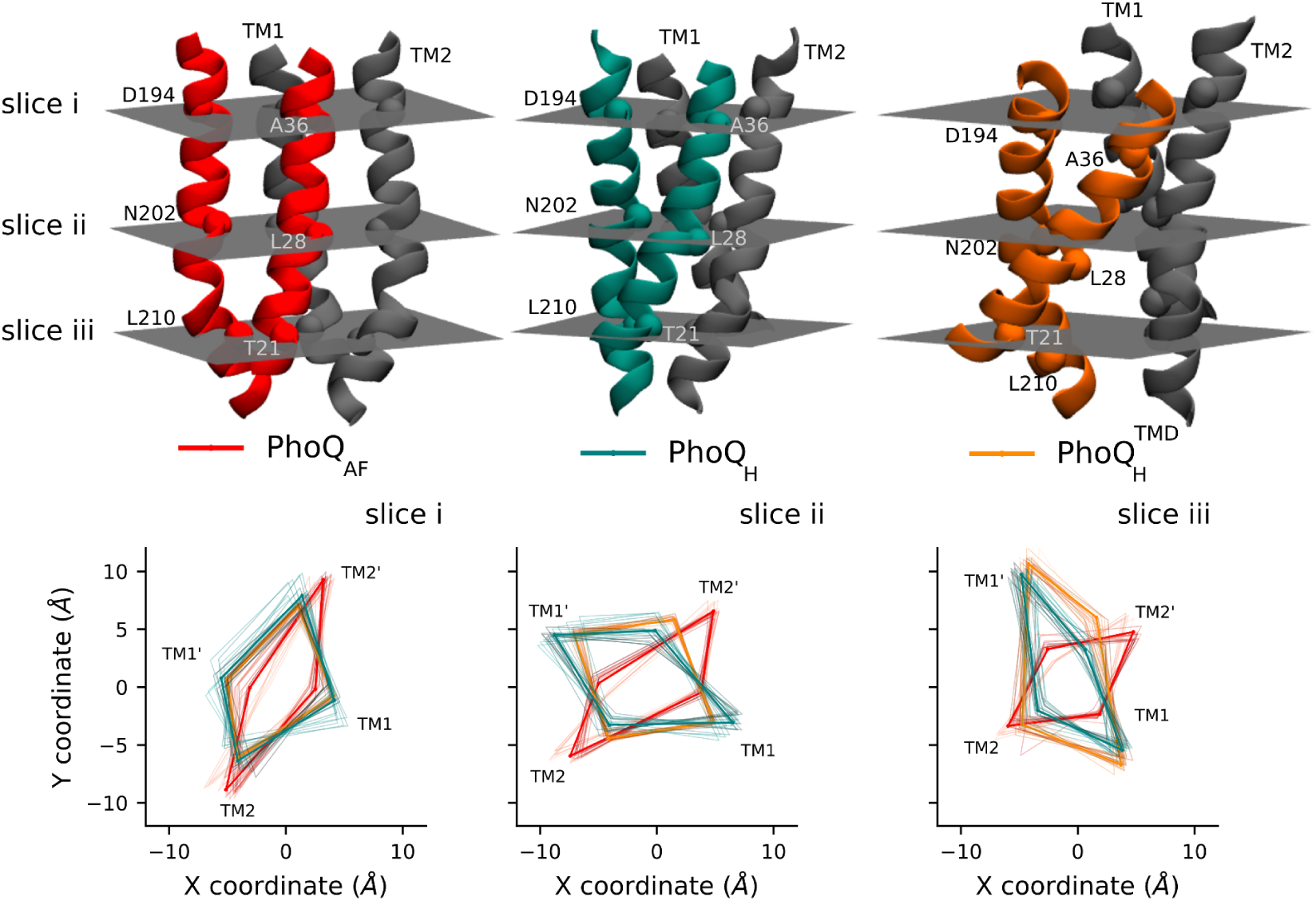
Interprotomer Cα Distances within TM Helices. Top panel: Cartoon representation of the PhoQ transmembrane (TM) helices divided into three slices (i-iii) defined by Cα atoms: (i) D194 and A36, (ii) N202 and L28, (iii) L210 and T21. Bottom panel: Distances measured during the MD simulations between Cα atoms within each slice for the *PhoQ^TMD^_AF_* (red), *PhoQ_H_* 𝐻 (teal), and *PhoQ^TMD^_AF_* (orange) models.

### Comparison to experimental cross-linking data

When compared to available experimental cross-linking data for the sensor interface, transmembrane (TM), and HAMP domains [38], all structural models exhibited generally good agreement between the cross-linking fractions and average inter-protomeric Cα-Cα distances (Figure 4). However, some key discrepancies emerged. Notably, the absence of cross-links for residues A20-L30 and the decreased periodicity for residues S43-D45 align with the larger interatomic distances and kink in the TM1 helix as it exits the membrane observed within the *PhoQ^TMD^_H_* model.

**Figure 4:**
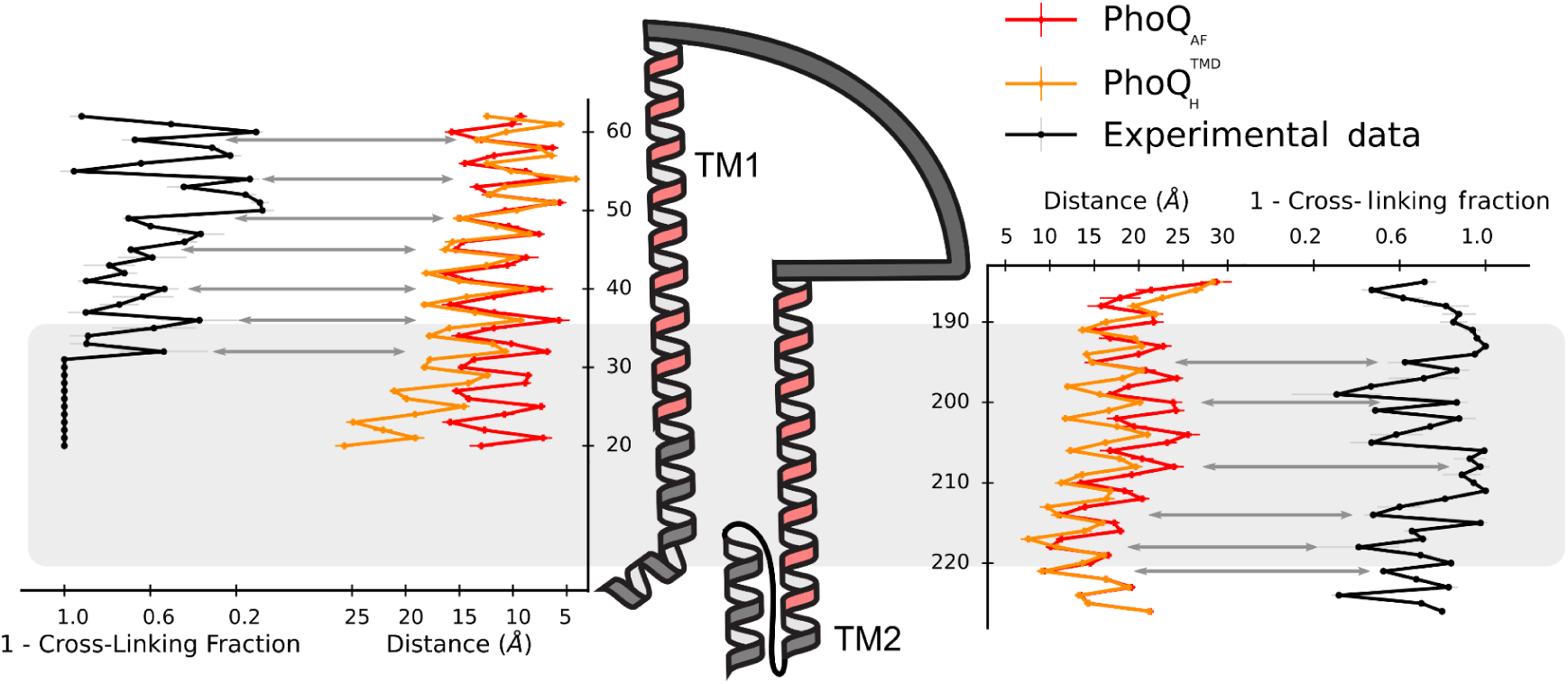
Comparison of cross-linking data with Cα distances in PhoQ models. Cross-linking data is represented in black, and the corresponding Cα inter-residue distances for *PhoQ_AF_* and *PhoQ^TMD^_H_*𝐻 are shown in red and orange, respectively. The membrane is depicted with a grey rectangle.

### Solvation of the transmembrane bundle

Previous experiments have shown that the polarity of N202 is essential for the function of PhoQ [37], [39]. This residue, located at the center of the transmembrane bundle, is thought to be crucial for solvating the bundle and enabling transitions between PhoQ signaling states. Consistent with this hypothesis, the *PhoQ^TMD^_H_* and *PhoQ_H_* models exhibited extensive transmembrane solvation throughout the simulations, contrasting with the complete absence of water molecules in the *PhoQ_AF_* model (Figure 5). Detailed analysis revealed a dynamic pattern of water occupancy within the transmembrane bundle. On average, 1.7 water molecules were present within the TM bundle of *PhoQ^TMD^_H_*. The most frequent configuration involved two water molecules occupying the bundle (45 ± 11 %). Occasionally, only one water molecule remained (23 ± 26 %), and rarely, the bundle became dehydrated (14 ± 12 %). The average residence time for an individual water molecule was around 287 ± 156 ns, with frequent exchanges occurring. Interestingly, some water molecules resided within the bundle for significantly longer durations, up to 1 microsecond (μs).

**Figure 5:**
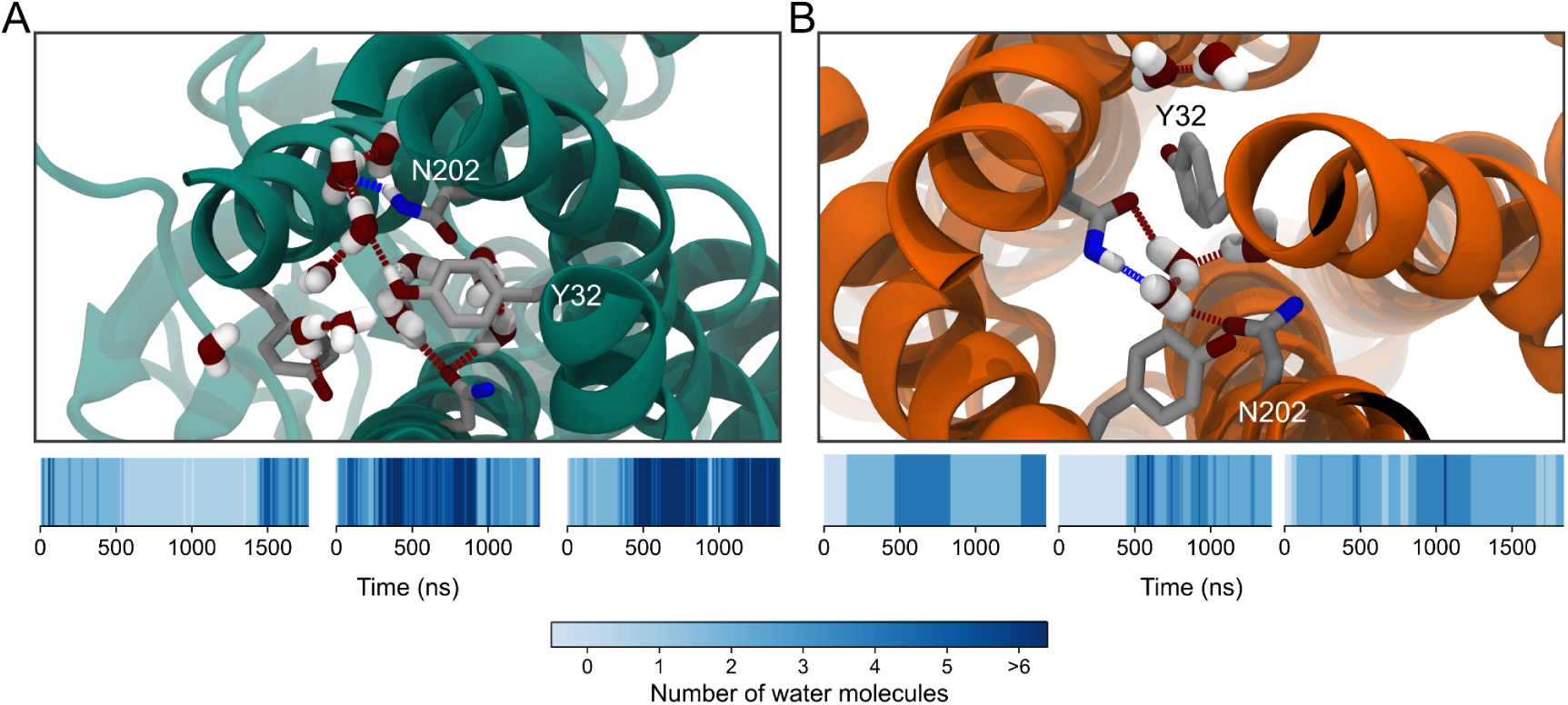
Solvation of the transmembrane domain. (A) and (B) Molecular representations of frames with the highest water occupancy for *PhoQ_H_* and *PhoQ^TMD^_H_* respectively. The protein is shown with a cartoon representation in teal (*PhoQ_H_*) and orange (*PhoQ^TMD^_H_*) and residues N202 and Y32 are highlighted in licorice. Water molecules are shown in red (oxygen) and white (hydrogen). The bottom blue intensity plots represent water molecule count throughout the simulations (three replicas), with higher intensity indicating more water molecules.

The transmembrane domain of *PhoQ_H_* displayed a greater degree of water occupancy, with an average of 3.5 water molecules per frame. The primary configurations involved either one (20 ± 30%) or two water molecules (19 ± 8 %) or a more dynamic distribution of three to seven water molecules (27 ± 24 %). The average water residence time was about 70 ± 50 ns, with one instance exhibiting a residence time of 1.4 μs.

### Interactions with Mg^2+^ cations

Considering the crucial role of PhoQ in Mg²⁺ sensing, we investigated the presence of Mg^2+^ ions in the vicinity of the acidic patch (residues: L131 - D150), known to be important for sensing cations. Minimal interactions with Mg^2+^ were observed for the *PhoQ_AF_* and *PhoQ^H^_AF_* models (less than 1% of the total simulation time for both chains). In striking contrast, the acidic patch of the *PhoQ^TMD^_H_* model exhibited extensive Mg²⁺ interactions with the acidic patch, particularly residues D149 and D152 of chain A (100% simulation time). Interestingly, each bound Mg²⁺ cation also interacted with a lipid phosphate group of the membrane, either directly or indirectly through water molecules (Figure 6). Although positioned further from the membrane surface, residue D125 in the acidic patch of chain B for *PhoQ^TMD^_H_* still bound to a Mg²⁺ ion for 95% of the simulation time.

**Figure 6:**
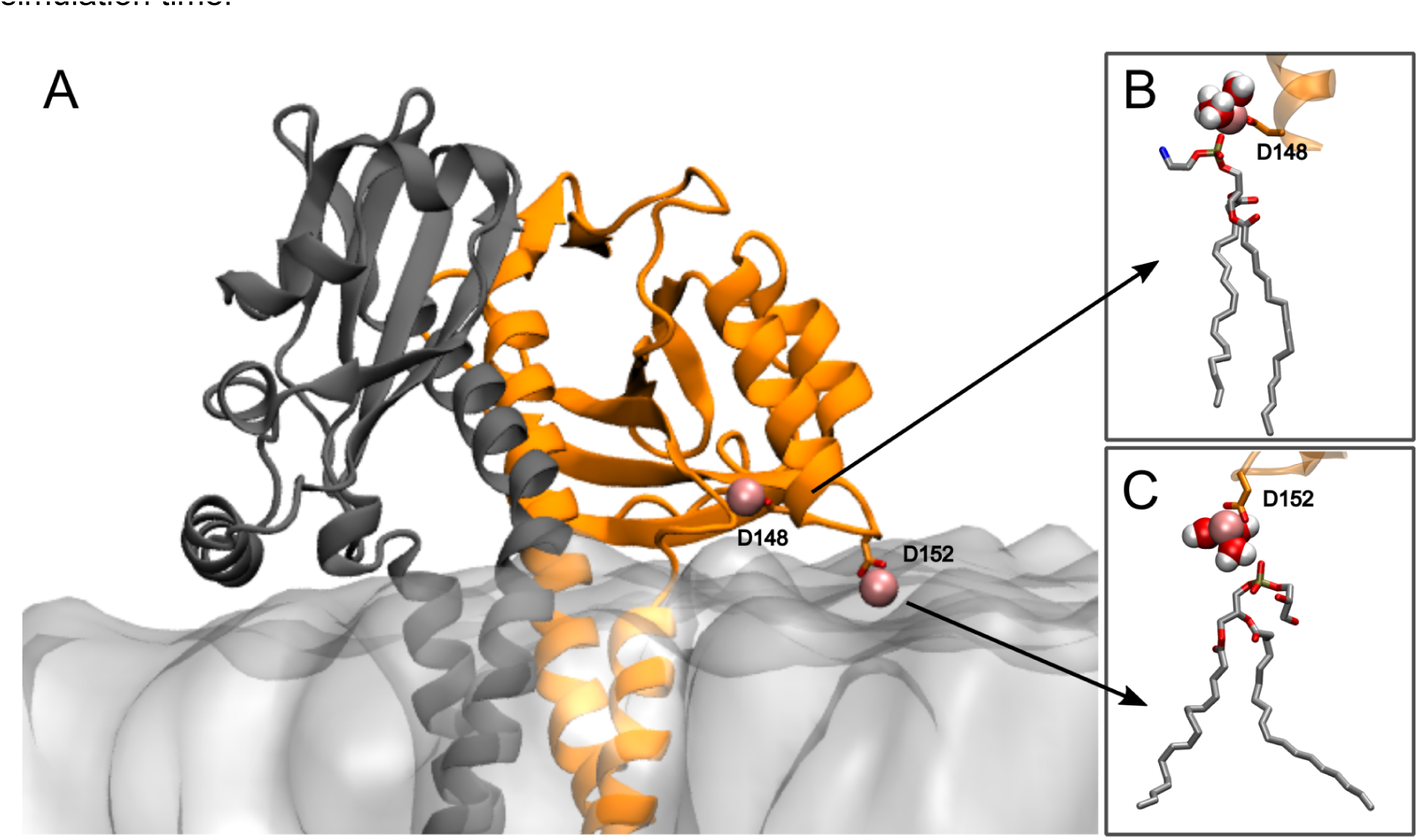
Mg²⁺-bridged interactions of the acidic patch and the membrane. (A) Cartoon representation of the sensor domain (gray/orange) interacting with the membrane (transparent gray surface). Mg²⁺ ions (pink spheres) bridge the acidic patch residues (D148, D152, sticks) and the membrane. Representative snapshots of interactions occurring directly (B) and via a water molecule (C).

### Dynamics of the cytoplasmic region

Despite overall similar appearances across simulations, principal component analysis (PCA) identified subtle but distinct conformational changes within the cytoplasmic domain of PhoQ (Supplementary Figure S10). The first principal component (PC1) captured a dominant bending motion in the B-chain of the S249-E296 helix (Supplementary Figure S10B, D), whereas the second principal component (PC2) described a scissoring motion of the N-terminal helices of the HAMP domain (R219-H234) (Supplementary Figure S10C, E). These small conformational shifts within the HAMP and DHp domains resulted in the dissociation of the catalytic domain from the core region in several simulations. To characterize these diverse conformations, we employed two key metrics: (i) the distance between a catalytic residue (H277) and the center of mass of the catalytic domain (excluding the lid region), and (ii) the angle formed between the Dhp domain core and the catalytic domain’s gripper helix (G445-E457). This analysis identified seven distinct poses adopted by the catalytic domain (Figure 7). The most prevalent conformation (red in Figure 7) closely resembled the Alphafold prediction, with the gripper helix bound to the DHp domain. Notably, at least one cytoplasmic domain adopted this conformation in almost all simulations (Supplementary Figure S11). Two additional conformations (light orange and green in Figure 7) exhibited an orientation that resembled a trans-phosphorylation state. These conformations positioned the catalytic domain near the core, with the gripper helix perpendicular to the Dhp domain. However, they crucially lacked interaction with the key residue H277. The light orange conformation, observed primarily in chain A of the *PhoQ^TMD^_H_* 𝐻 model, positions the domain slightly closer to the core compared to the green conformation. The remaining clusters (yellow, purple, cyan, and orange) all represented detached conformations of the catalytic domain lacking interaction with the Dhp core. Notably, the cyan cluster corresponded to a conformation where the domain moved towards and directly interacted with the membrane surface.

**Figure 7:**
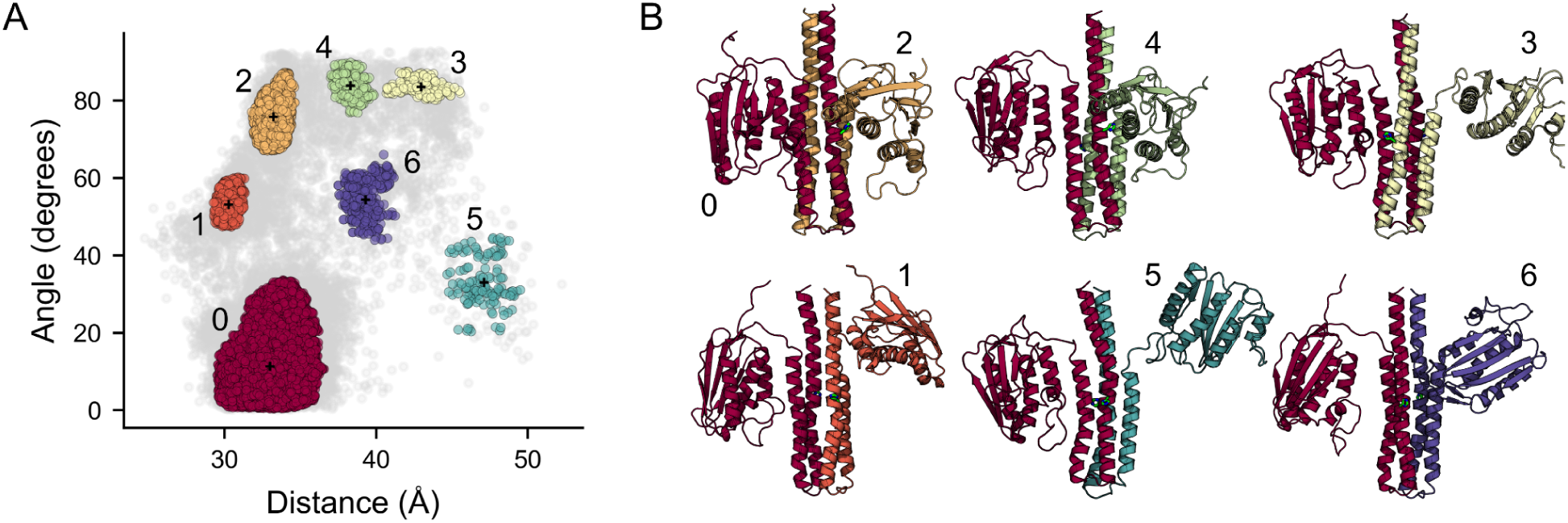
Conformational dynamics of the Catalytic Domain. A) Scatter plot depicting the distance between H277 (DHp bundle) and the catalytic domain’s center of mass (x-axis) versus the angle between the gripper helix and the DHp core axis (y-axis). Each chain was analyzed separately and the data was merged in the plot. A multi-step clustering identified seven clusters (colored dots) and gray dots represent transition conformations between clusters. The black cross indicates the center of mass of each cluster. B) Cartoon representations of the major identified conformations. Each chain is colored according to its cluster assignment in panel (A).

## Discussion

This study leveraged AlphaFold structural predictions and MD simulations to explore the conformational dynamics of PhoQ, a sensor protein critical for bacterial signal transduction. Although the initial AlphaFold2 prediction displayed high confidence and domain arrangements consistent with expectations, it depicted a near-parallel interface in the sensor domain, unlike the ∼40° angle observed in the known X-ray structure (PDB ID: 3bq8). By integrating a previously published model of the transmembrane domain [39] with targeted MD simulations, we successfully captured two additional conformations of PhoQ (hybrid models: *PhoQ^TMD^_H_* and *PhoQ_H_*). Even though all models remained stable and compatible with available cross-linking data [38], small structural deviations within the sensor and transmembrane domains resulted in substantial overall conformational changes between the models, providing insights into PhoQ signaling at atomic resolution (Figure 8, Supplementary Movie S3).

**Figure 8:**
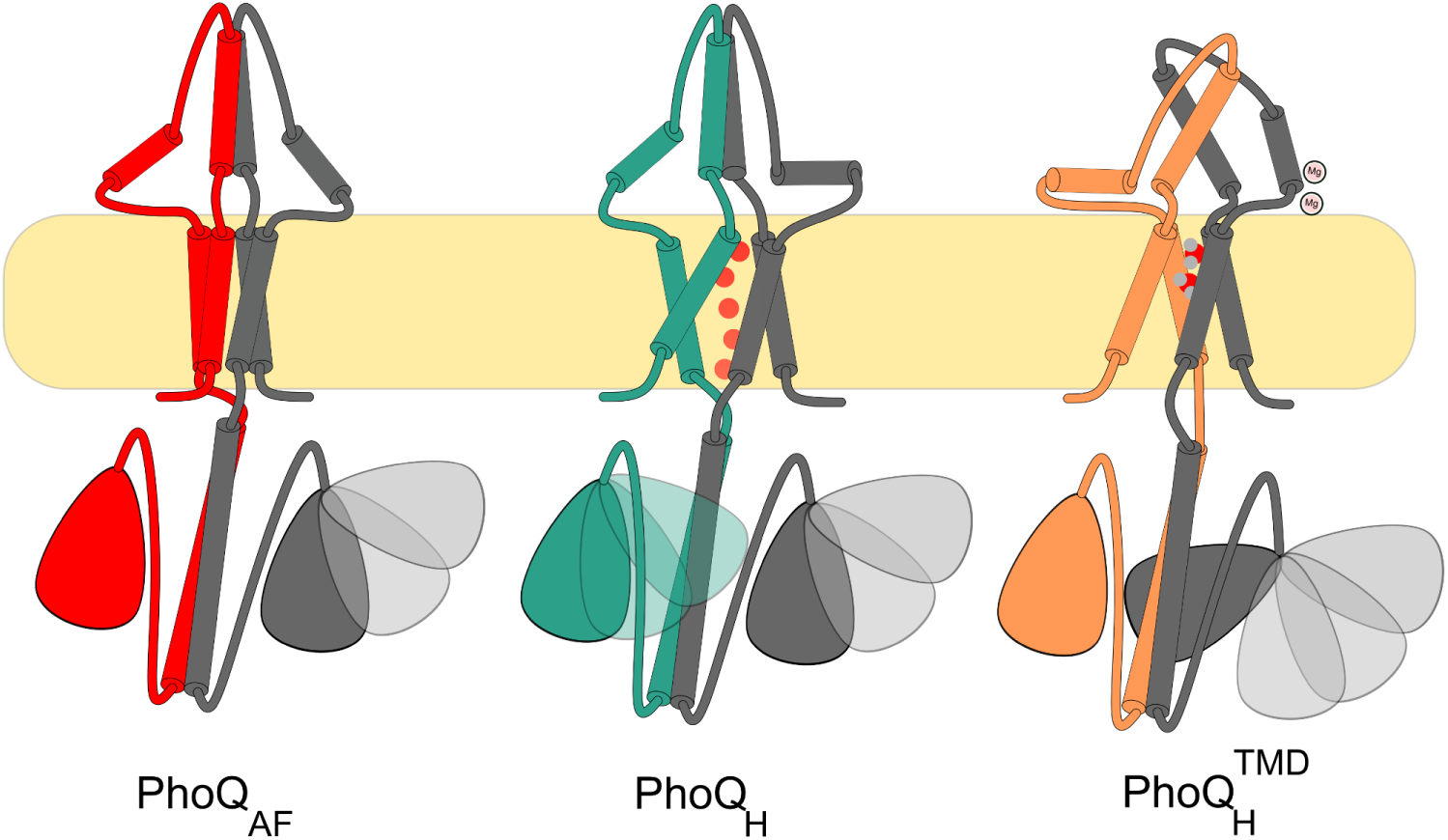
Schematic representation of identified conformations of PhoQ within its signaling landscape. Three potential conformations of the PhoQ signaling protein were identified through computational modeling. AlphaFold predicted a stable and symmetrical structure (*PhoQ_AF_*, red), which might represent a pre-activated state before autophosphorylation. A hydrated conformation (*PhoQ_H_*, teal) could correspond to a transition state during signaling. Finally, a highly asymmetric structure with minimal hydration and two bound Mg²⁺ ions (*PhoQ^TMD^_AH_*, orange) indicates the repressed state of PhoQ.

In particular, the hybrid structures exhibited a high degree of asymmetry, aligning with previous evidence suggesting an asymmetric nature of PhoQ signaling [13], [31], and generally of histidine kinases [33]. The opening of the sensor domain interface resulted in the reorientation of one of the acidic patches of *PhoQ^TMD^_H_*, thus allowing the bridging of Mg cation with the membrane bilayer. This observation corresponds perfectly with previous hypotheses based on X-ray structures [31], [48]. These studies proposed that orienting the acidic patch towards the membrane through Mg^2+^ bridges triggers PhoQ signaling. This bridging interaction between Mg^2+^ and both the acidic patch and the membrane is crucial since the negatively charged acidic patch and the negatively charged phosphate groups in the membrane would otherwise repel each other. The Mg^2+^ interaction occurred either directly with membrane phosphate groups or mediated through a water molecule. Based on this, we propose that *PhoQ^TMD^_H_* represents the repressed state of PhoQ, where Mg^2+^ binding stabilizes the opening of the sensor interface helices.

In contrast to the complete absence of water molecules observed in *PhoQ_AF_* and the partial hydration of *PhoQ^TMD^_H_*, *PhoQ_H_* displayed a significantly higher degree of water molecules occupying the core of its transmembrane domain. This finding agrees with prior research indicating the importance of the polar residue N202 for signaling and the proposed presence of a water-filled cavity [37], [39], also apparent in the HtrII structure [36], [58]. It was hypothesized that this hydration pocket facilitates transitions between the different states of PhoQ, by relaxing the tertiary packing interactions in the bundle [37]. The high level of hydration in *PhoQ_H_* TM could indicate that this conformation is an intermediate state within the signaling landscape of PhoQ. The higher intrinsic preference of the sensor domain for the kinase on-state (not bound to Mg^2+^) could explain the residual hydration observed in *PhoQ^TMD^_H_*, which would allow an easier switching away from the repressed state [37], [59].

Despite significant conformational changes in the sensor and transmembrane domains, the HAMP and DHp domains remained remarkably similar across all conformations. This observation lends further support to the previously proposed allosteric coupling model [60] between these domains, where communication occurs through subtle structural changes rather than a single, concerted conformational shift. Notably, a slight bend in the cytoplasmic S-helix, aligning with solved structures of similar domains [44], and a scissoring motion observed in the HAMP domain, consistent with proposed signaling of HAMP domain [61], [62], appear sufficient to transmit these signals towards the catalytic domain. Furthermore, the catalytic domain exhibited higher mobility in the *PhoQ^TMD^_H_* (repressed state) simulations. Although none of the observed catalytic domain conformations facilitated autophosphorylation of the histidine residue, this enhanced mobility could represent another regulatory mechanism for repressing PhoP phosphorylation.

Beyond elucidating PhoQ’s specific activation pathway, these insights hold significant promise for guiding future investigations into the broader histidine kinase family. The shared structural features observed in PhoQ suggest a potential for these findings to be extrapolated to related proteins, paving the way for a more comprehensive understanding of histidine kinase signaling mechanisms across various biological systems. Furthermore, the validity of our generated models opens exciting avenues for further studies on PhoQ itself. These models can be instrumental in exploring PhoQ’s modulation by small peptides [63] and antimicrobial agents, potentially leading to the development of novel therapeutic strategies.

## Methods

### Molecular modeling

The initial model was obtained using the sequence of *E. coli* K12 (P23837) and the colab implementation of AlphaFold2-multimer [47], with default parameters. The hybrid model was constructed by replacing the transmembrane domain of the predicted model with the transmembrane domain from a previously published atomic model [39]. Specifically, residues M1 to S43 in TM1 and S193 to L224 in TM2 were exchanged. To ensure smooth junctions between the different models, the Rosetta kinematic closure protocol (KIC) was applied for relaxation [64].

### Molecular Dynamics

The models of PhoQ dimers were embedded in a bacterial mimetic bilayer composed of a 3:1 ratio of phosphatidylethanolamine (POPE) and phosphatidylglycerol (POPG). The system was solvated with a water box with a padding of 20 Å and neutralized using NaCl (150 mM) and MgCl_2_ (5 mM).

The simulations were performed with the CHARMM36 force field [65], including CMAP corrections for the protein. The water molecules were described with the TIP3P water parameterization [66].

### Targeted Molecular Dynamics simulations

Targeted molecular dynamics simulations using NAMD [67] were used to enforce the conformation of the interface helices in the sensor domain observed in the X-ray structure (PDB: 3bq8). The cutoff for non-bonded interactions was set to 12 Å with a switching distance at 10 Å. Periodic electrostatic interactions were computed using Particle-Mesh Ewald (PME) summation with a grid spacing smaller than 1 Å. To maintain a constant temperature of 310 K, Langevin dynamics with a damping coefficient of 1.0 ps was employed. Additionally, a constant pressure of 1 atm was maintained using the Langevin barostat [68].

To preserve the stability of the secondary structure during the simulation, extra bonds were applied to the torsional angles of the protein’s backbone. The targeted dynamics simulation was carried out over 50 ns. The interface helices were then kept fixed for an additional 50 ns before finally releasing all constraints, except for the secondary structure, and allowing the system to evolve for an extra 50 ns.

### Unrestrained molecular dynamics

Each system underwent extended Molecular Dynamics simulations using the OpenMM molecular engine [69]. The systems were first minimized for 5000 steps followed by an equilibration (1.75 ns), progressively releasing positional restraints on the backbone atoms. The cutoff for non-bonded interactions was set to 12 Å with a switching distance at 10 Å. The periodic electrostatic interactions were computed using particle-mesh Ewald (PME) summation. Constant temperature of 310 K was imposed by Langevin dynamics with a damping coefficient of 1.0 ps. Constant pressure of 1 atm was maintained with Monte Carlo barostat [70]. The hydrogen mass repartitioning scheme was used to achieve a 4 fs time-step [71]. Snapshots from each simulation were extracted at 1 ns time intervals for structural analysis. Each simulation was carried out up to at least 1.4 us (Supplementary Table S2).

### Analyses

We employed a combination of custom scripting and established software packages to analyze the data. Custom Tcl scripts were written to automate the extraction of features within VMD software [72], [73], [74]. The extracted data was analyzed with pandas [75], [76] and NumPy [77], and graphics were generated using Matplotlib [78]. The Principal Component Analysis (PCA) was carried out on the alpha carbon using the implementation provided by the ProDy library [79]. The curvature of the helices was measured with the Bendix tool [80]. The visualization and molecular renderings were produced with VMD and Pymol [81], [82]. Morphing was performed using ChimeraX [83], [84], [85].

## Acknowledgments

T.L. acknowledges funding from Swiss National Science Foundation (SNSF: PCEFP3_194606) and J.Y. from the Max Planck Society.

## Author contributions

S.L. study design, data acquisition, data analysis, article drafting, critical revision, and final approval of the manuscript. J.Y. conception of the idea, study design, article drafting, critical revision, and final approval of the manuscript. T.L. conception of the idea, study design, article drafting, critical revision, and final approval of the manuscript.

## Data availability statement

All simulations are available at 10.5281/zenodo.10988292

Script for analyses are available at https://github.com/symelaz/PhoQ

## Additional Information

The authors declare no conflicts of interest.

