## Supplementary Material for "Atomic insights into the signaling landscape of *E. coli* PhoQ Histidine Kinase from Molecular Dynamics simulations"

Symela Lazaridi: 0009-0003-3323-7215

Jing Yuan: 0000-0003-1219-3316

Thomas Lemmin: 0000-0001-5705-4964

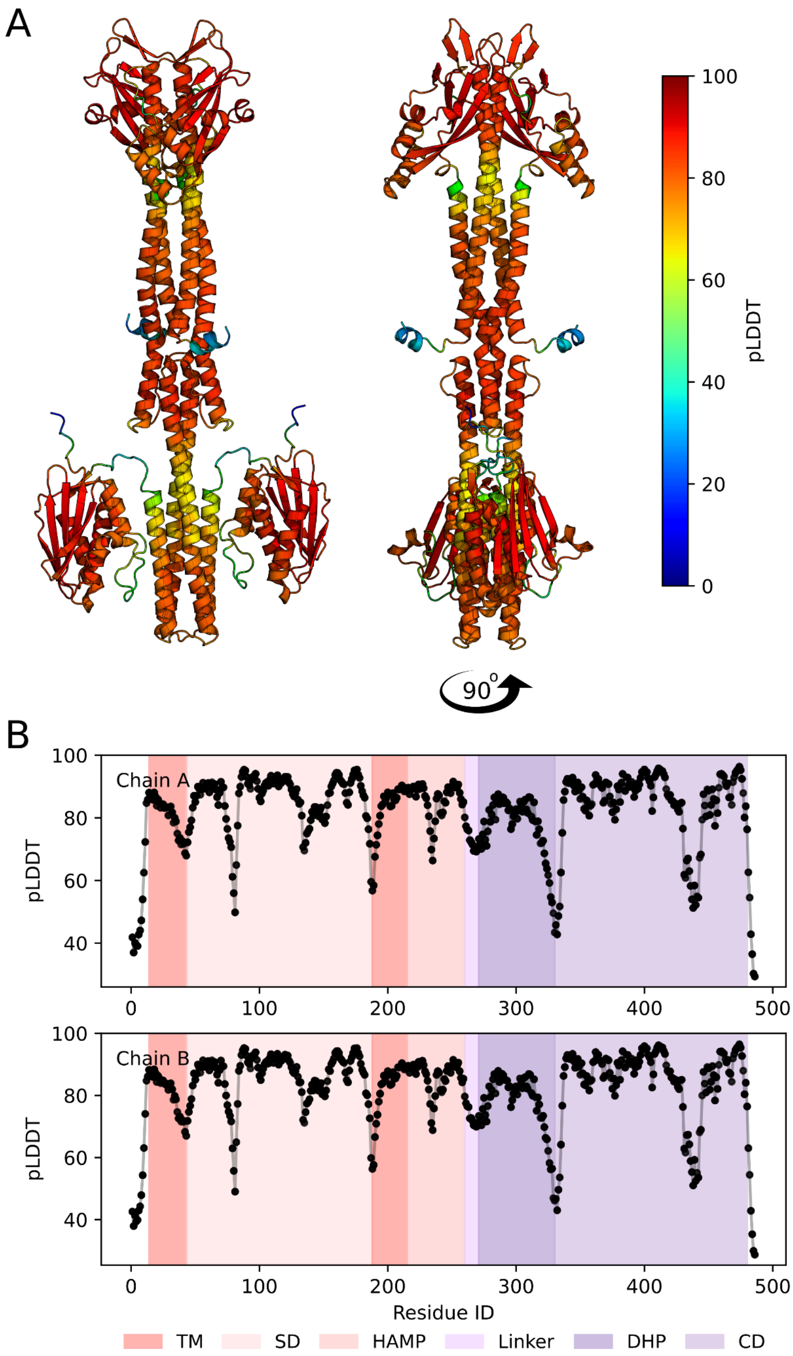


**Figure S1: pLDDT score of AlphaFold model.** A) Cartoon representation of the PhoQ protein model generated by AlphaFold-multimer. The coloring scheme indicates pLDDT score, ranging from blue (low confidence) to red (high confidence). B) Plot of pLDDT scores across all amino acid positions in the PhoQ protein sequence. The different domains (sensor, transmembrane, HAMP, Linker, DHp, and catalytic) are highlighted with shades of red and purple.

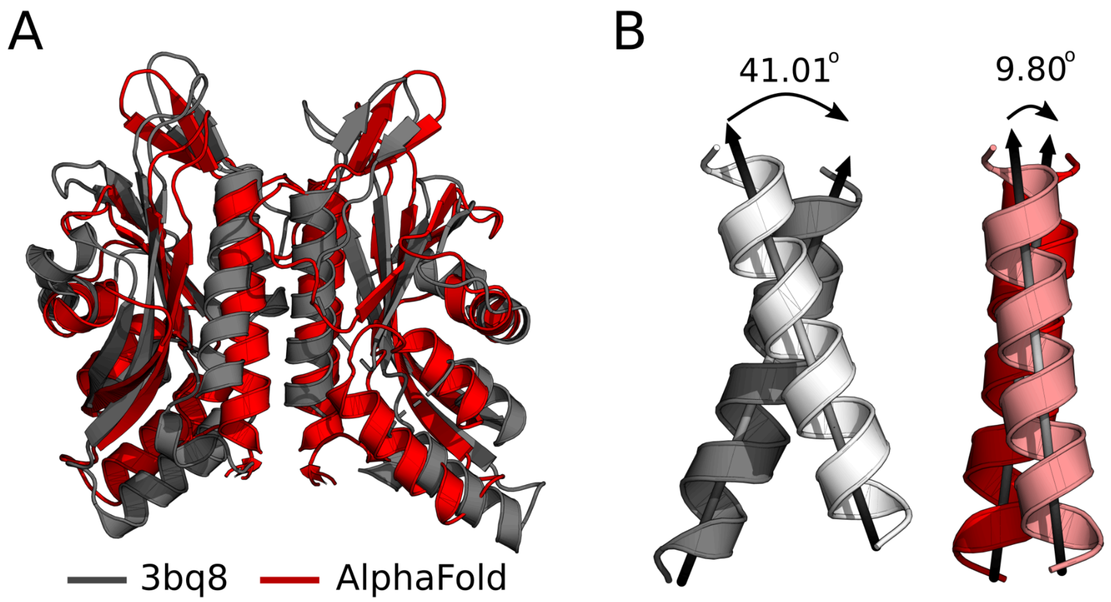


**Figure S2: Comparative analysis of the AlphaFold sensor domain with the 3bq8 structure.** A) The sensor domain of the AlphaFold model (red) aligned with the experimentally solved structure (PDB id: 3bq8) (gray). B) Visualization of interhelical angles at the sensor interface. The experimental structure (gray and white) is compared to the AlphaFold prediction (red and pink).

**Table S1: Structural search of the AlphaFold predicted model against the Protein DataBank (PDB).**

| Model | Domain | Hits for chain A | | | Hits for chain B | | |
|---|---|---|---|---|---|---|---|
| | | PDB id | RMSD (Å) | TM-Score | PDB id | RMSD(Å) | TM-Score |
| AlphaFold predicted Structure | SD | 1yax_C | 1.37 | 0.94 | 1yax_C | 1.36 | 0.94 |
| | TM | No hits found | | | No hits found | | |
| | HAMP | 2y21_J | 2.27 | 0.61 | 2y21_J | 2.35 | 0.62 |
| | DHp | 5uky_B | 1.18 | 0.84 | 5uky_B | 1.19 | 0.84 |
| | CD | 1id0_A | 1.7 | 0.89 | 1id0_A | 1.71 | 0.89 |

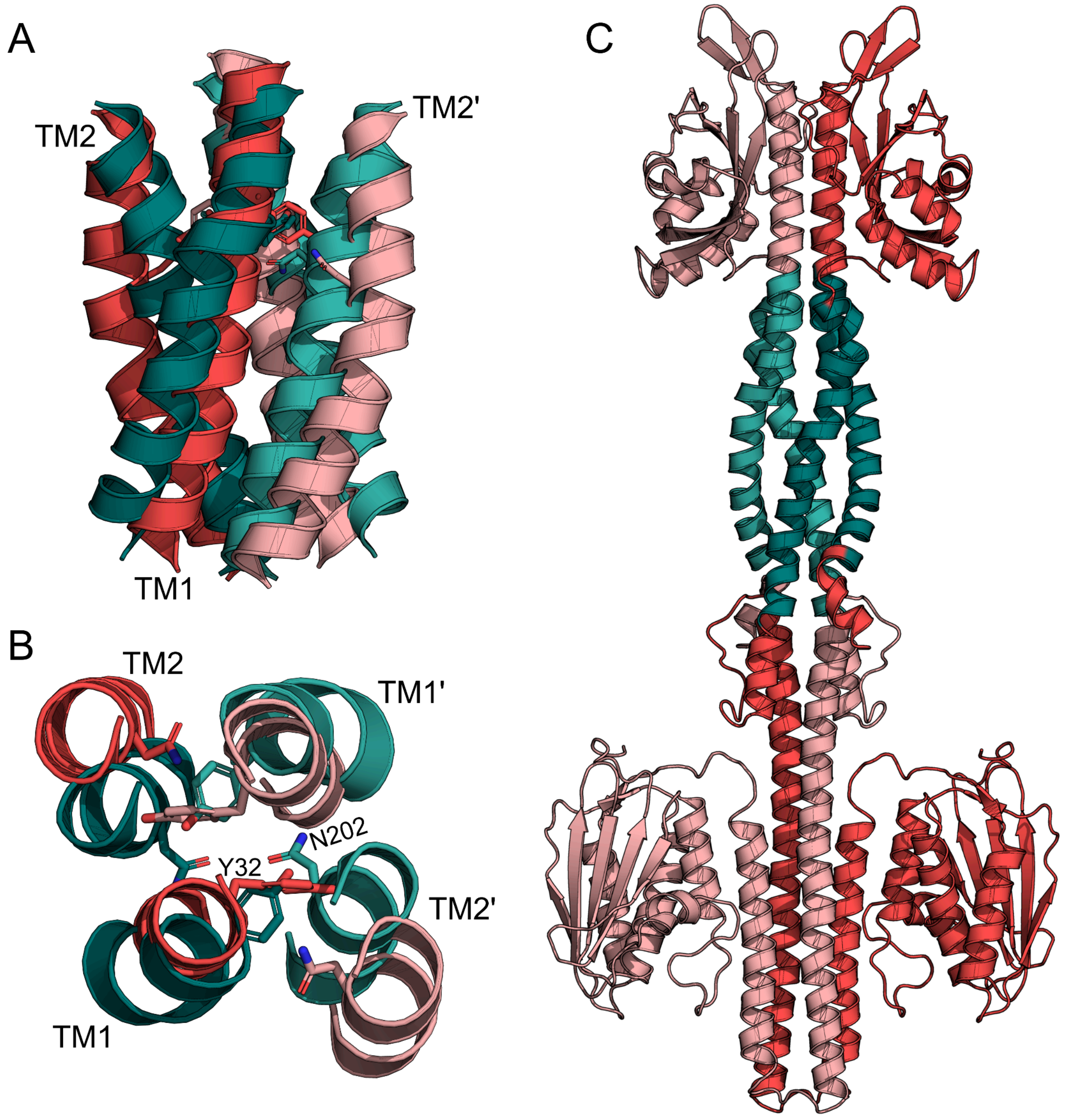


**Figure S3: Comparison of transmembrane domains**
Cartoon representation of the transmembrane domains viewed from the periplasmic side (A) and through the membrane (B). The AlphaFold predicted and modeled transmembrane domaines are colored in red and in teals, respectively. C) Full length PhoQ structure after exchanging the transmembrane domain with the modeled structure.

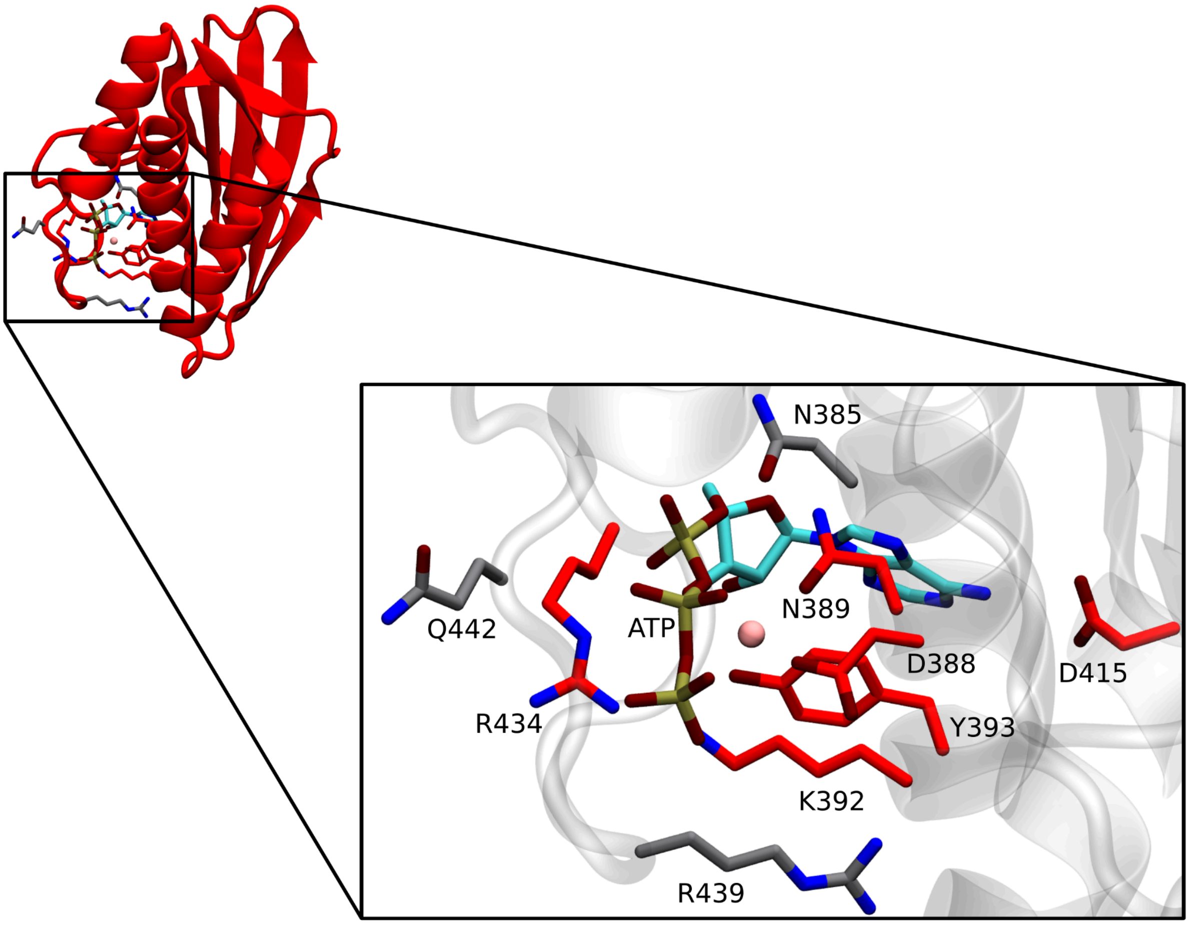


**Figure S4: Visualization of an ATP molecule docked within the pocket of the AlphaFold model.** The ATP molecule is depicted in a stick representation, with carbon atoms colored cyan, oxygen atoms in red, and phosphorus atoms in tan. The catalytic domain is represented as a cartoon. Residues interacting with ATP in the experimentally solved structure (PDB id 1IDO) are visualized as sticks, highlighted in red if they are also predicted to interact by the AlphaFold model, and in gray, if not.

**Table S2: Summary of simulations**

Simulation durations are reported in ns, the average and standard deviation of the RMSD are reported in Ångstrom for both the full protein and the core region (core: M1-E486), excluding the catalytic domain.

| | $PhoQ_{AF}$ | | | $PhoQ_{AF}^{TMD}$ | | | $PhoQ_{H}$ | | | $PhoQ_{H}^{TMD}$ | | |
|---|---|---|---|---|---|---|---|---|---|---|---|---|
| | Frames | RMSD (Å) | | Frames | RMSD (Å) | | Frames | RMSD (Å) | | Frames | RMSD (Å) | |
| | | Full | Core | | Full | Core | | Full | Core | | Full | Core |
| **Replica 1** | **1363** | **6.83 $\pm$ 0.87** | **3.62 $\pm$ 0.44** | **1476** | **10.32 $\pm$ 3.76** | **5.19 $\pm$ 2.03** | **1770** | **12.75 $\pm$ 3.53** | **4.16 $\pm$ 0.47** | **1461** | **7.79 $\pm$ 1.63** | **5.13 $\pm$ 1.14** |
| **Replica 2** | **1440** | **11.17 $\pm$ 2.33** | **5.94 $\pm$ 1.30** | | **-** | **-** | **1342** | **6.03 $\pm$ 0.93** | **4.56 $\pm$ 0.79** | **1404** | **5.54 $\pm$ 1.06** | **4.00 $\pm$ 0.77** |
| **Replica 3** | **1368** | **8.82 $\pm$ 0.93** | **3.43 $\pm$ 0.47** | | **-** | **-** | **1399** | **7.08 $\pm$ 1.18** | **5.15 $\pm$ 0.94** | **1836** | **9.03 $\pm$ 4.63** | **4.19 $\pm$ 0.89** |
| **Total** | **4171** | **8.76 $\pm$ 2.57** | **4.23 $\pm$ 1.28** | **1476** | **10.32 $\pm$ 3.76** | **5.19 $\pm$ 2.03** | **4511** | **9.06 $\pm$ 3.81** | **4.60 $\pm$ 0.82** | **4701** | **7.46 $\pm$ 3.24** | **4.54 $\pm$ 1.00** |

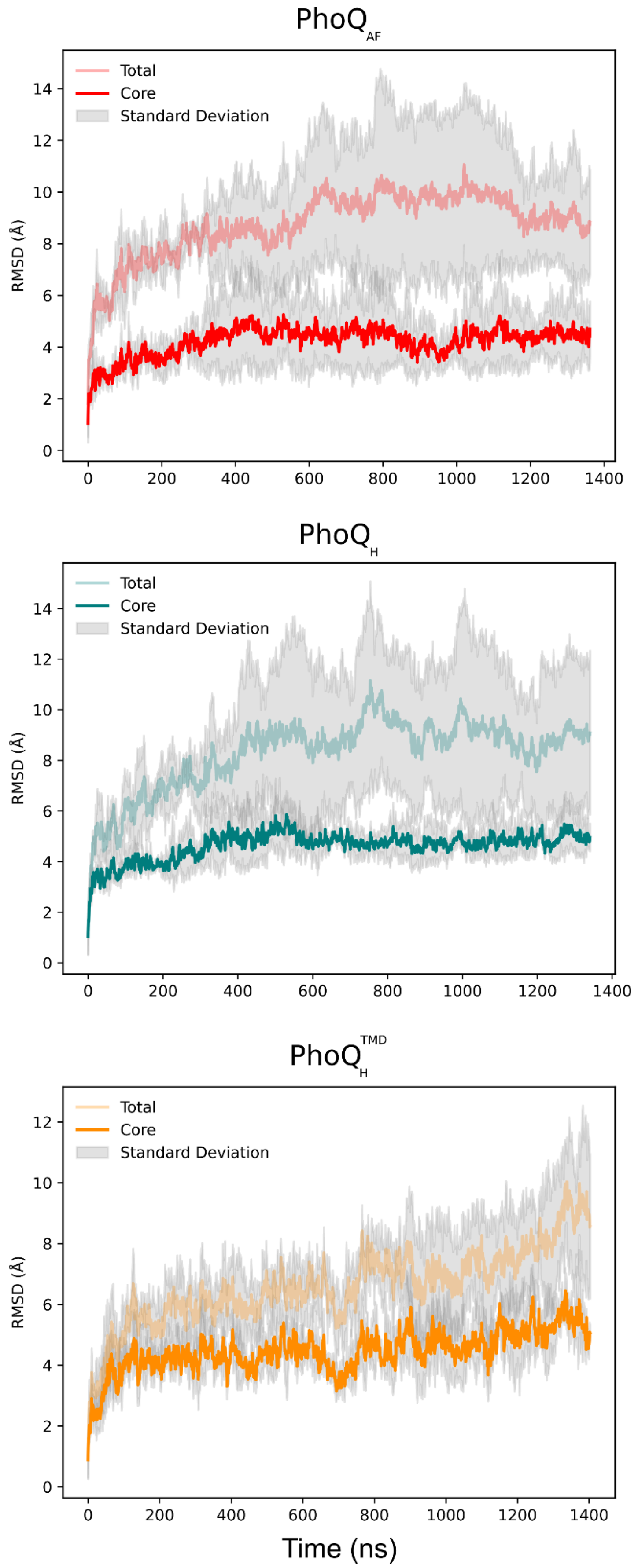


**Figure S5: RMSD of PhoQ models across simulations.** The mean RMSD calculated over all three replicates is shown for $PhoQ_{AF}$ (red), $PhoQ_{H}$ (teal), and $PhoQ_{H}^{TMD}$ (orange). Standard deviation is depicted in gray. Deeper color shades represent the RMSD of the PhoQ core (residues M1-G330), excluding catalytic domains, while lighter shades represent the full protein RMSD.

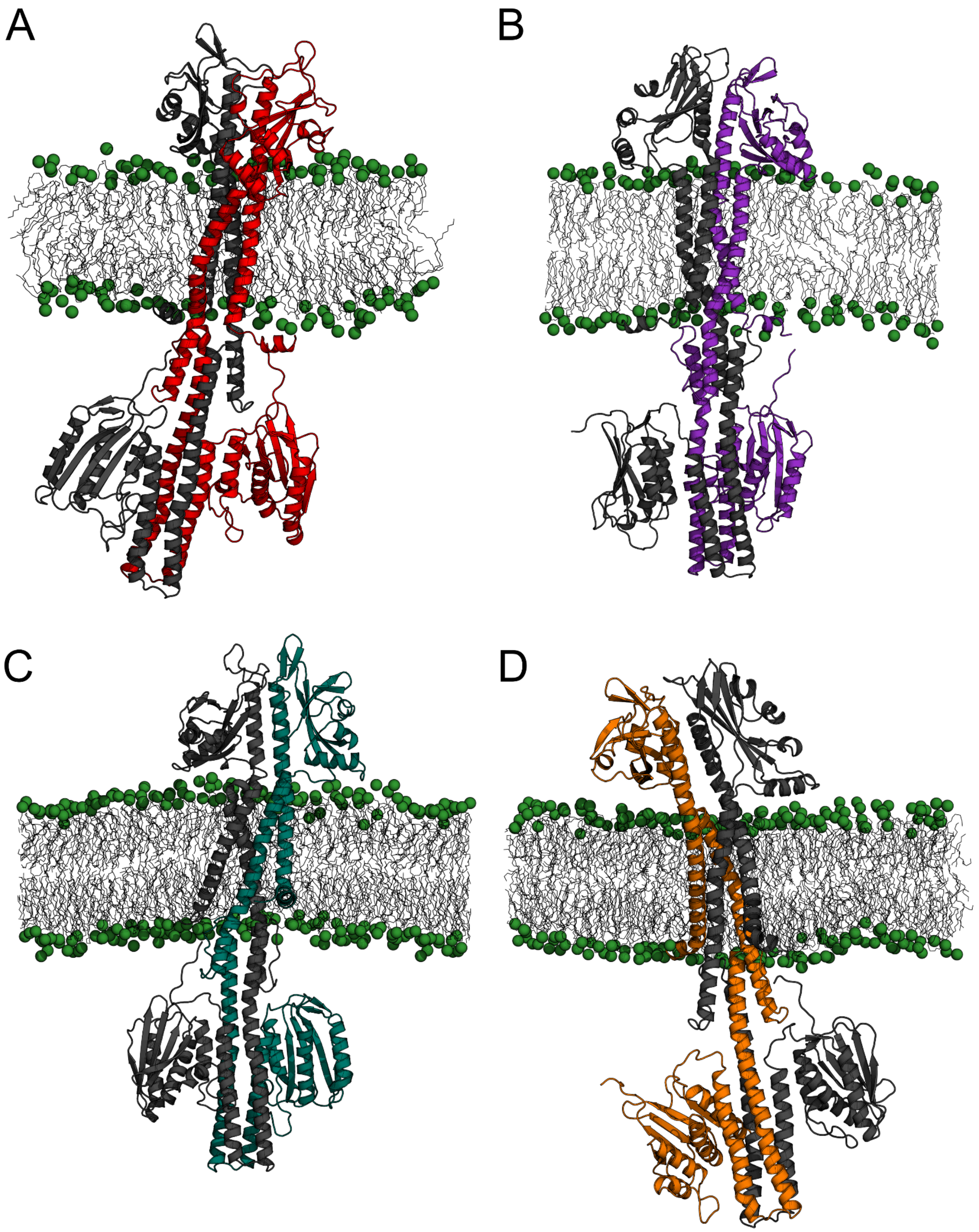


**Figure S6: Representative final structures of PhoQ models from molecular dynamics simulations.** Panels depict cartoon representations of A) $PhoQ_{AF}$ model, B) $PhoQ_{AF}^{TMD}$, C) $PhoQ_{H}$ and D) $PhoQ_{H}^{TMD}$. Different protein chains are colored in gray and red, purple, teal, or orange, respectively. The hydrophobic core of the membrane is shown with gray sticks, and phosphate groups are depicted as green spheres.

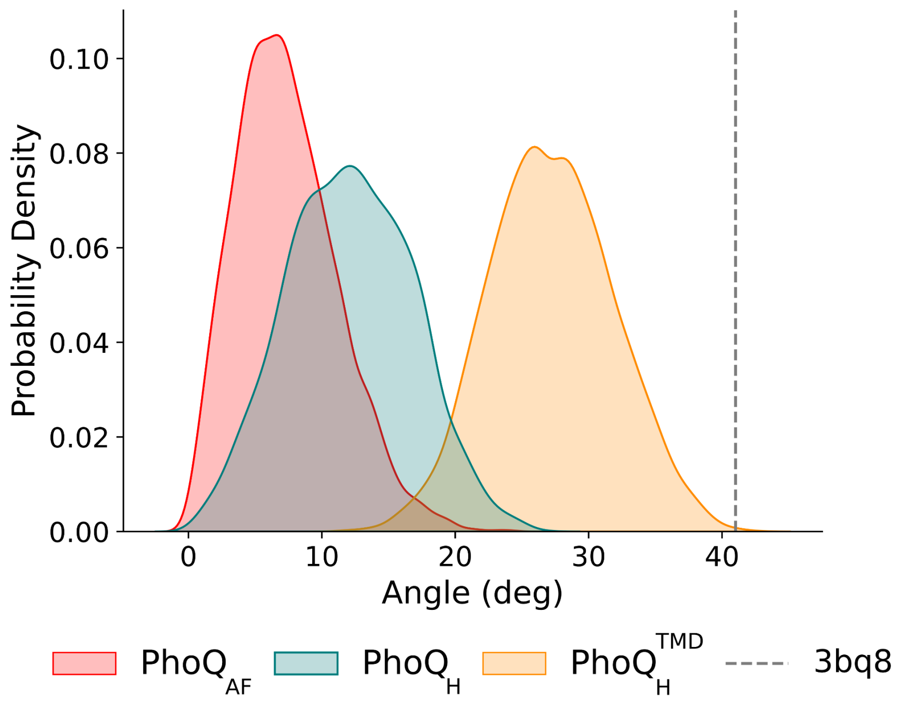


**Figure S7: Distribution of PhoQ sensor interface cross-angles.** Kernel density estimation (KDE) of the sensor interface cross-angle measured during molecular dynamics simulations for $PhoQ_{AF}$ (in red), $PhoQ_{H}$ (in teal) and $PhoQ_{H}^{TMD}$ (in orange). The vertical dashed line indicates the cross-angle observed in the experimentally solved structure (PDB ID: 3bq8).

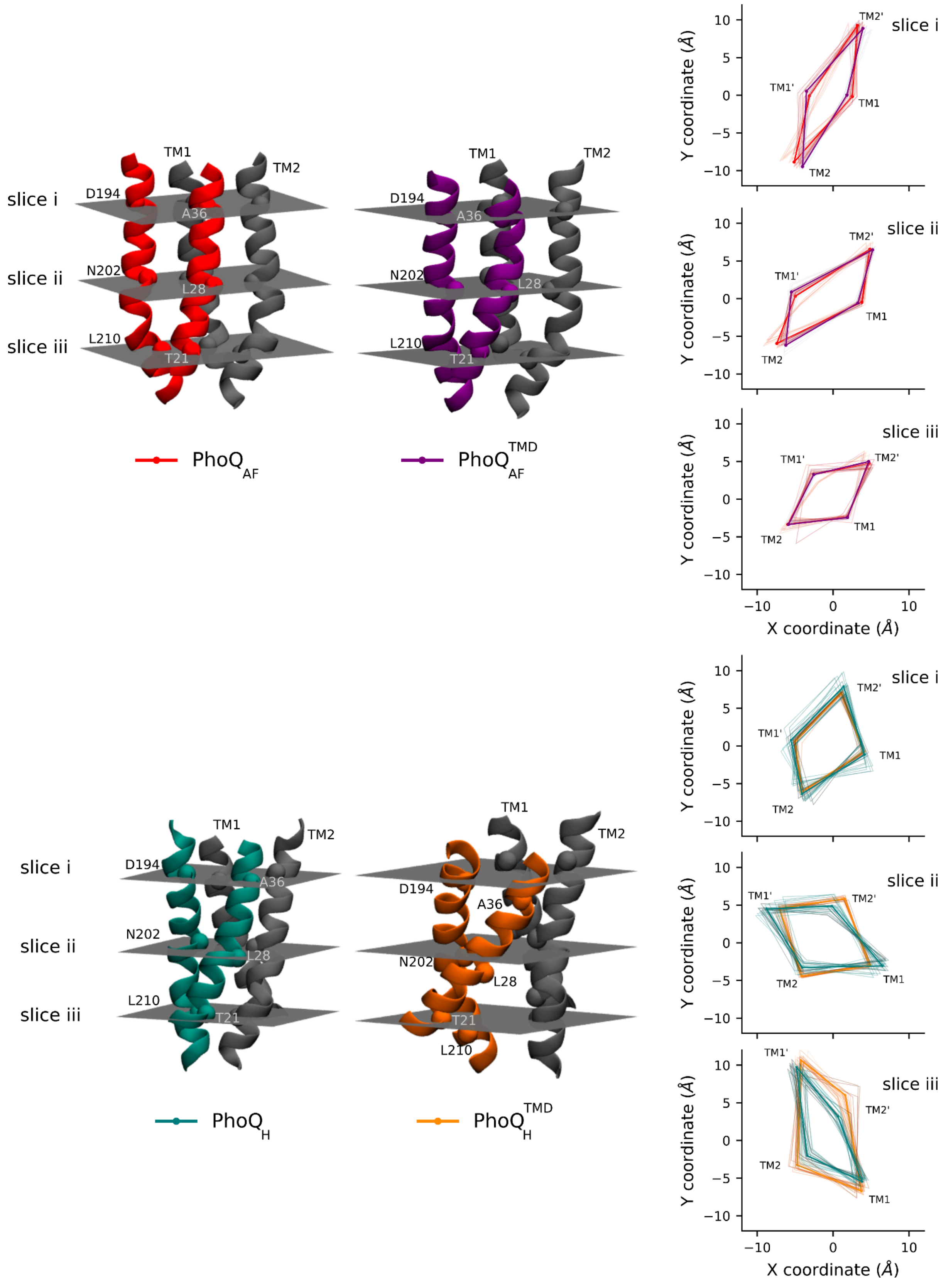


**Figure S8: Comparison of the interprotomer distances in the transmembrane bundle**
Left panel: Cartoon representation of the PhoQ transmembrane (TM) helices divided into three slices (i-iii) defined by Cα atoms: (i) ASP194 and ALA36, (ii) ASN202 and LEU28, and (iii) LEU210 and THR21. Bottom panel: Distances measured during the MD simulations between Cα atoms within each slice for the $PhoQ_{AF}$ (red), $PhoQ_{AF}^{TMD}$ (purple), $PhoQ_{H}$ (teal) and $PhoQ_{H}^{TMD}$ (orange).

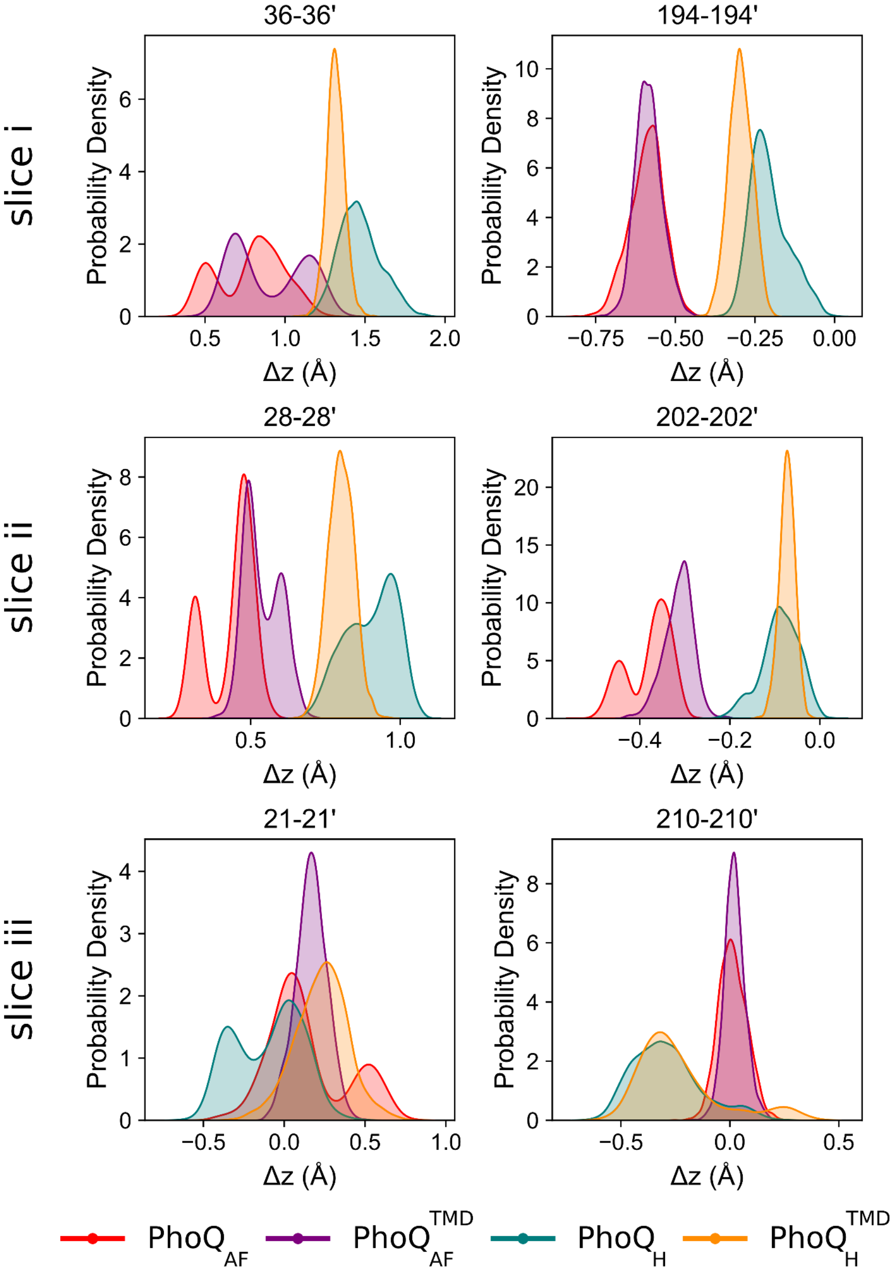


**Figure S9: Pistoning motion of the transmembrane bundle**
Kernel density estimation (KDE) of displacement along the z-axis between pair of residues belonging the slices defined in Figure S8: $PhoQ_{AF}$ (red), $PhoQ_{AF}^{TMD}$ (purple), $PhoQ_{H}$ (teal) and $PhoQ_{H}^{TMD}$ (orange).

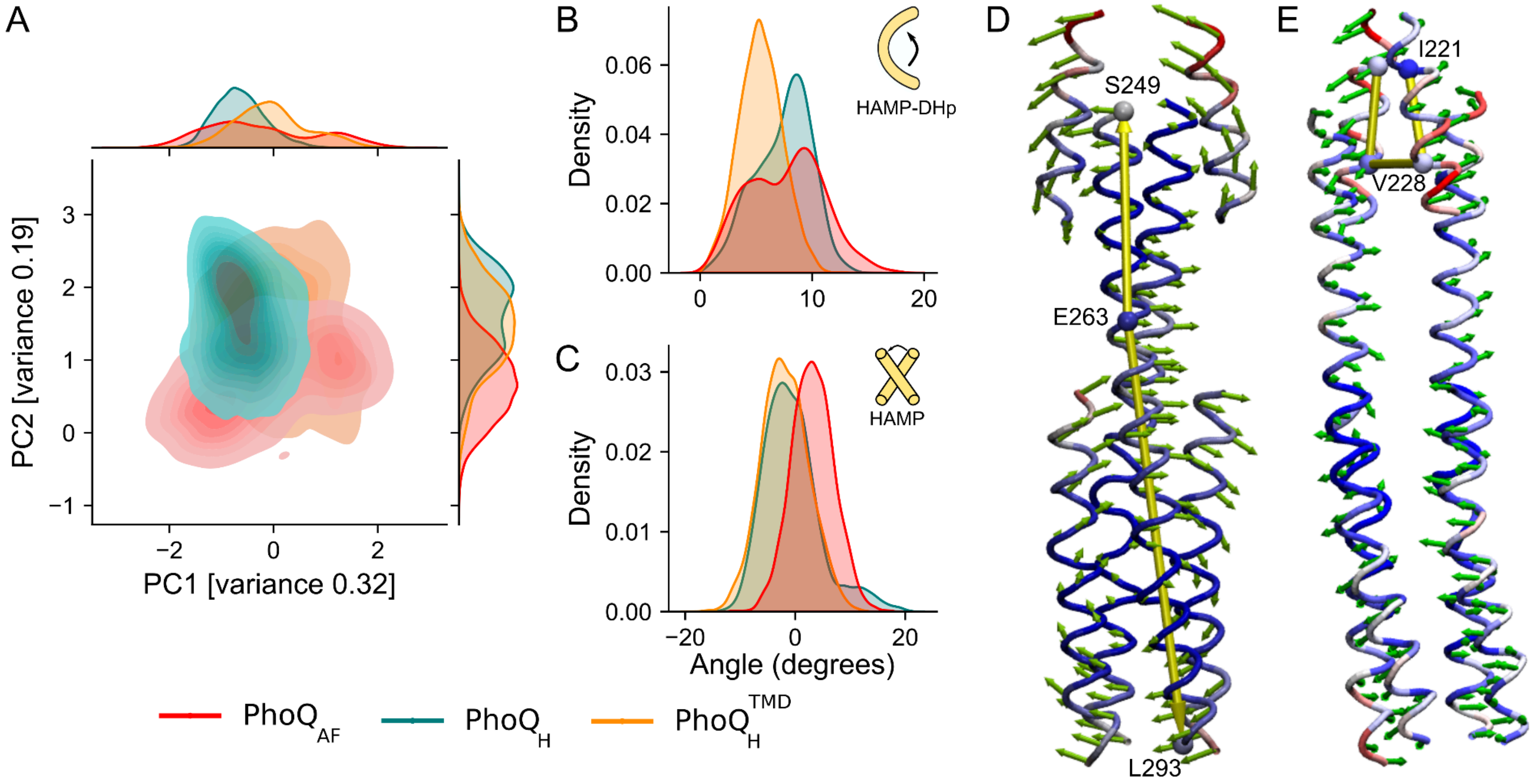


**Figure S10: Principal Component Analysis (PCA) of the cytoplasmic domains**

A) Projection of the conformations of $PhoQ_{AF}$ (red), $PhoQ_{H}$ (teal) and $PhoQ_{H}^{TMD}$ (orange) into the eigen space defined by the first two principal components. PCA was computed using the backbone atoms of the structured regions within the HAMP, S-helix, and Dhp domains. Marginal distributions for each PC are displayed on the respective axes. B) Distribution of the curvature of the S249 to L293 measured at position E263. representative of PC1. C) distribution of the torsional angle of the HAMP domain defined between. I221–V228 and V228' – I221', representative of PC2. Eigenvectors corresponding to PC1 (D) and PC2 (E) observed during the simulations are represented as green arrows. The models are coloured to reflect the magnitude of the displacement, from small (blue) to larger movement (red). Animation of the PCs is reported in Movies S1 and S2.

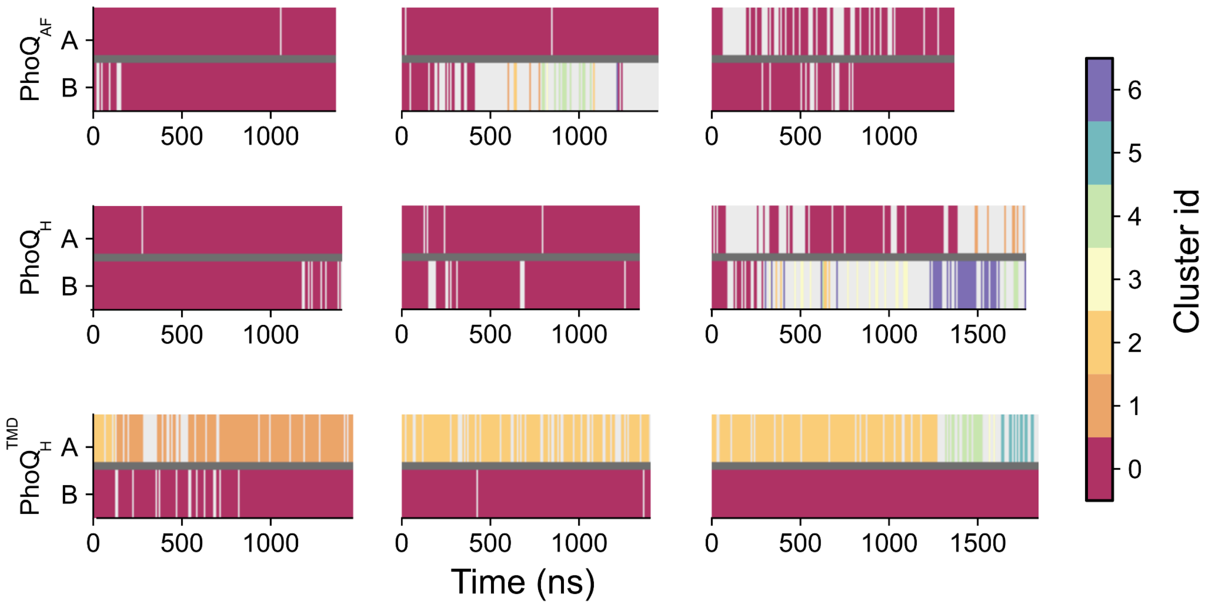


**Figure S11: Conformational dynamics of the catalytic domain**
Timeline showing the conformations of catalytic structures sampled during the molecular dynamics simulation for each chain of PhoQ$_{AF}$, PhoQ$_{H}$, PhoQ$_{H}^{TMD}$. Each stripe of the timeline is colored according to the clusters defined in Figure7.

**Movie S1: Animation PC1:** Figure S10D
**Movie S2: Animation PC2:** Figure S10E

**Movie S3: Activation and deactivation of PhoQ:**
The movie illustrates the shift between PhoQ's extreme states, beginning with the active state (PhoQ$_{AF}$), transitioning to the partially active state (PhoQ$_{H}^{TMD}$), and returning back to the active state.
